# Unbiased and comprehensive identification of viral encoded circular RNAs in a large range of viral species and families

**DOI:** 10.1101/2024.06.24.600382

**Authors:** Alexis S Chasseur, Maxime Bellefroid, Mathilde Galais, Meijiao Gong, Sarah Mathieu, Camille Ponsard, Laure Vreux, Carlo Yague-Sanz, Benjamin G Dewals, Nicolas A Gillet, Benoît Muylkens, Carine Van Lint, Damien Coupeau

**Author notes:** Corresponding author : Carine Van Lint. Université libre de Bruxelles (ULB); Service of Molecular Virology; Rue Profs Jeener & Brachet, 12; 6041 Gosselies; Belgium. Ragon Institute of MGH, MIT and Harvard, Cambridge, MA 02139, USA. Department of Medical Oncology, Jerome Lipper Multiple Myeloma Center, Dana-Farber Cancer Institute and Harvard Medical School, Boston, MA, USA. Author Contributions: A.S.C., B.M., D.C. wrote the manuscript; B.G.D., B.M., C.V.L., D.C. designed the research project; A.S.C., C.Y.S., D.C. designed vCircTrappist; A.S.C., M.B., M.G., M.G., S.M., C.P., L.V. conducted the research and processed the samples; B.G.D., N.A.G., B.M., C.V.L., D.C. supplied the reagents and samples; B.G.D., B.M., C.V.L. earned the fundings; A.S.C., B.M., D.C. analyzed the data; M.B., M.G., M.G., S.M., C.P., L.V., C.Y.S., B.G.D., N.A.G., C.V.L. reviewed the manuscript; D.C. is the leader of the project. **Competing Interest Statement:** The authors declare no conflict of interests in the present study.

## Abstract

Non-coding RNAs play a significant role in viral infection cycles, with recent attention focused on circular RNAs (circRNAs) originating from various viral families. Notably, these circRNAs have been associated with oncogenesis and alterations in viral fitness. However, identifying their expression has proven more challenging than initially anticipated due to unique viral characteristics. This challenge has the potential to impede progress in our understanding of viral circRNAs. Key hurdles in working with viral genomes include: (1) the presence of repetitive regions that can lead to misalignment of sequencing reads, and (2) unconventional splicing mechanisms that deviate from conserved eukaryotic patterns.

To address these challenges, we developed vCircTrappist, a bioinformatic pipeline tailored to identify backsplicing events and pinpoint loci expressing circRNAs in RNA sequencing data. Applying this pipeline, we obtained novel insights from both new and existing datasets encompassing a range of animal and human pathogens belonging to Herpesviridae, Retroviridae, Adenoviridae and Orthomyxoviridae families. Subsequent RT-PCR and Sanger sequencings validated the accuracy of the developed bioinformatic tool for a selection of new candidate viral encoded circRNAs. These findings demonstrate that vCircTrappist is an open and unbiased approach for comprehensive identification of virus-derived circRNAs.

**Significance Statement:** Circular RNAs (circRNAs) were revealed to have prominent roles in cellular life in the past decade. They were more recently shown to be expressed by viruses, influencing their infectious cycles and host-pathogen relationship. In this context, viruses that were not previously associated with cellular splicing processes are shown to express circRNAs through unknown mechanisms. These non-canonical circRNAs were already shown to be important in the viral cycle and pathogenesis of the viruses they are encoded from. Here, we propose a bioinformatics pipeline that bypasses the limitations of the existing tools in the identification of viral circRNA. Using this pipeline, we discovered numerous candidates and invite the reader to start its own exploration in the realm of viral encoded circRNAs.

**Graphical Abstract:** 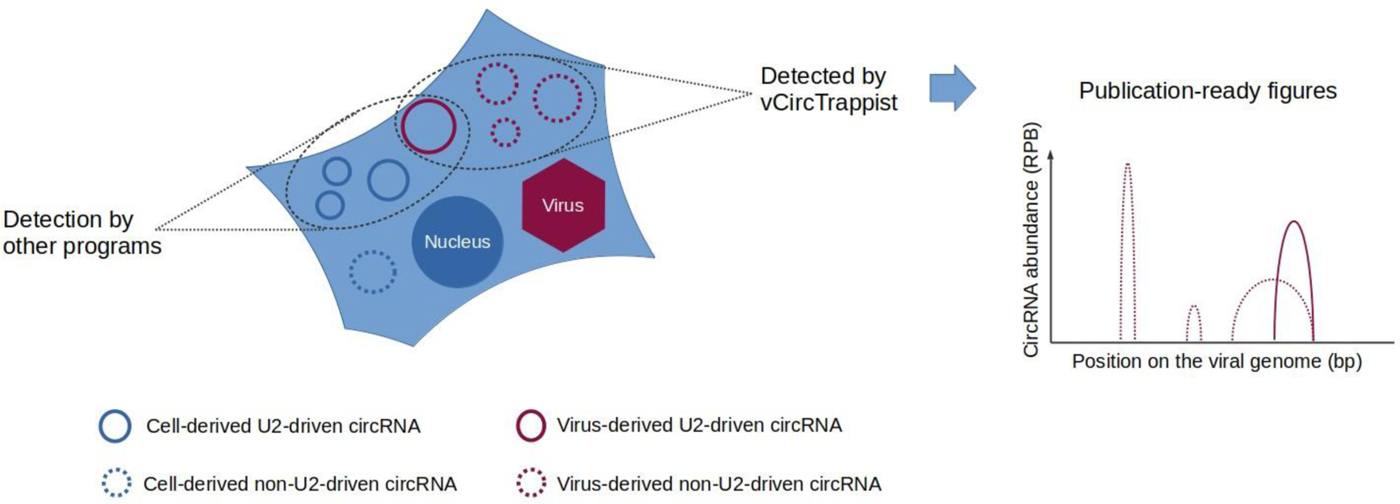

## Introduction

While not classified as living organisms, viruses show remarkable adaptability to their environment. They exhibit an exceptional ability to co-opt and manipulate various cellular pathways, using the cellular machinery for their regulation, replication and dissemination. Viruses perturb every facet of the central dogma of biology, spanning from genomic (1) to proteomic levels (2).

Circular RNAs (circRNAs) have been implicated in diverse molecular mechanisms. Initially considered non-coding, they have since been demonstrated to undergo translation, akin to conventional linear mRNAs. In the field of viral biology, circRNAs have been identified across numerous viral families (reviewed in (3–5)). While the majority of their functions remain to be elucidated, it is worth emphasizing that their diversity and multiple splicing patterns involved in their biogenesis render their individual study challenging (6). Nonetheless, in-depth characterization of select circRNA candidates within specific viral families has provided promising insights into their potential roles in pathogenesis. One notable example is circE7 from Human Papillomavirus (HPV) 16, which has been demonstrated to undergo translation, encoding the oncoprotein E7 through an RNA methylation-mediated mechanism (7). Furthermore, roles in pathogenesis have been uncovered for herpesviruses (8, 9), influenzavirus (IAV) H1N1 (10), and Hepatitis C virus (HCV) (11).

Biogenesis of viral circRNAs presents an intriguing puzzle. While the generation of cellular circRNAs was mostly associated with a canonical backsplicing mechanism involving the U2 splicing machinery, viral circRNAs appear to be partly produced through as-yet-unknown mechanisms. This phenomenon has been observed primarily in RNA viruses (10, 12–15), but it is also noteworthy in DNA viruses, exemplified by Marek’s Disease Virus (MDV, also known as Gallid Herpesvirus 2 – GaHV-2) (16), which displays numerous circRNAs with splicing patterns not associated with the canonical GU-AG motif, surrounding upstream and downstream ends of removed introns. It is noteworthy the GU-AG backsplicing signature, associated with the U2 machinery, is a pattern in circRNA identification by most existing bioinformatics tools. However, this signature is not representative of the diversity of circRNA junction observed in the viral world. For instance, in the case of the Respiratory syncytial virus (RSV) (14), an AU-UA pattern was observed, but a clear association with any splicing mechanism remains elusive. A recent study exploring HCV circRNAs (11) has suggested that the generation of functionally active viral circRNAs might result from an alternative mechanism due to the absence of viral RNA within the nucleus, where canonical splicing occurs. In the case of the coronavirus Murine Hepatitis Virus (MHV), a study from Yang and colleagues has underscored the importance of the viral exonuclease NSP14 in circRNA biogenesis (17).

To decipher the intricacies of the viral infectious cycle and its impact on host cellular processes, it is imperative to comprehensively catalog all viral factors expressed during infection. This identification phase presents technical challenges. Within this context, the study of viral circRNAs is marked by pronounced practical difficulties. Indeed, viral circRNAs might be produced by various mechanisms and viral genomes are characterized by the presence of numerous repeated regions, these two features making circRNA identification more challenging in viruses. While a lot of programs are already available to identify new circRNA candidates (reviewed in (18)), only three are commonly used in the context of viral infections : CIRI2 (19), find_circ (20) and circRNA_finder (21). Using these, a database was created by Fu and collaborators and named VirusCircBase (22). However, these currently available tools all use the fixed GU/AG signal to identify circRNAs. Their second limitation is the fact that they do not take into account the peculiarities of viral genomes described above. In the present study, we introduce vCircTrappist, a novel bioinformatics pipeline tailored to the distinctive attributes of viral genomes, thereby surmounting the constraints associated with prevailing circRNA analysis tools. vCircTrappist uses data generated by Illumina high-throughput sequencing to precisely detect atypical splicing events, called backsplice junctions (BSJ), enabling the identification of uncharted circRNA candidates. Stringent filters, designed to exclude potential artifacts stemming from viral features such as genomic repetitive elements, are integrated to curtail false positives. Moreover, vCircTrappist is engineered to discern circRNAs exhibiting non-canonical splicing patterns from the canonical U2 ones. Leveraging this innovative framework, and by using newly and pre-existing datasets, we have successfully uncovered circRNAs expressed by diverse viral families, some of which were experimentally validated. This pioneering program promises to enrich resources like VirusCircBase (22) with a plethora of circRNA candidates, thereby substantially elevating the discovery rate of significant viral circRNAs.

## Materials and Methods

### Cell lines and viruses

The Embryonic Stem Cell-Derived Line 1 (ESCDL-1) was kindly given to us by Caroline Denesvre. It was cultured, as indicated in Vautherot *et al* (23), at 37°C – 5% CO2 in Dubelcco’s Modified Eagle Medium (DMEM) F12 (32500035 – Gibco) supplemented in 1% non-essential amino acids (11140050 – Gibco), 1mM sodium pyruvate (11360070 – Gibco), 50U/mL of penicillin and streptomycin (15070063 – Gibco) and 10% fetal bovine serum (FBS) (10270106 – Gibco). They were infected with the very virulent strain of MDV, RB-1B. In this case, the cells were first transfected using a viral GFP bacmid where the internal repeat long is lacking, as described in (24). One million cells were transfected using the Amaxa fibroblast kit (VPI-1002 – Lonza) and the program F-024, following the manufacturer’s protocol. After that, the infection was propagated 3 times to reach around 80% of infected cells in a culture of three million cells. We removed bioinformatically the internal repeats in the sequence for the subsequent analysis.

The two productive bovine leukemia virus (BLV)-infected ovine cell lines L267_LTaxSN_ (1) and YR2_LTaxSN_ (25) used in this study constitutively express the viral transactivator TaxBLV as a result of transduction with the pLTaxSN retroviral vector of native L267 (26) or YR2 (27, 28) cells, respectively. The ovine cell lines L267 and YR2 were established from the L267 B-cell lymphoma and M395 B-cell leukemia, respectively, developed by BLV-infected sheeps (accession number : KT122858.1) (29). All BLV-infected cell lines were maintained in Opti-MEM medium (31985070 – Gibco) supplemented with 10% FBS (10270106 – Gibco), 1 mM sodium pyruvate (11360070 – Gibco), 2 mM L-glutamine (25030149 – Gibco), 1% non-essential amino acids (11140050 – Gibco) and 100 μg/ml kanamycin monosulphate (KAN0025 – Formedium).

The 293T-BLV samples were obtained by calcium-phosphate transfection (CalPhos Mammalian Transfection Kit; 631312 – TaKaRa) of 293T cells (CRL-3216 – ATCC) with the pBLV344 plasmid containing a complete and infectious proviral copy of BLV (29). Fourty-eight hours post-transfection, RNA was extracted as described below. 293T cells were maintained in DMEM (11965092 – Gibco) supplemented with 10% FBS (10270106 – Gibco), 1 mM sodium pyruvate (11360070 – Gibco), and 50U/mL of penicillin and streptomycin (15070063 – Gibco).

The human T lymphotropic virus 1 (HTLV-1) productively-infected SLB1 (RRID: CVCL_RT63) and HUT-102 (RRID: CVCL_3526) cell lines were cultured in RPMI 1640 medium (52400-025 – Gibco) supplemented with 50 U/ml of penicillin and streptomycin (15070063 – Gibco), with 20% FBS for the SLB1 cell line and 10% FBS (10270106 – Gibco) for the HUT-102 cell line.

The human lung carcinoma cell line A549 (CCL-185 – ATCC) was cultured in DMEM (11965092 – Gibco) supplemented with 10% FBS (10270106 – Gibco). These cells were infected with the human Adenovirus C5 (hAdV C5) (accession number : AC_000008.1 ; VR-5 – ATCC) as described in (30). Lymphoblastoid cell lines (LCLs) were derived from calves developing malignant catarrhal fever (MCF) upon infection with strain C500 of alcelaphine gammaherpesvirus 1 (AlHV-1 ; accession number : KX905135.1) (31, 32). The virus was propagated in bovine turbinate (BT) fibroblasts (CRL-1390 – ATCC) and maintained by limited passage (<5) before calf infection.

For the supplemental data and figures, we also explored different viral strains, namely: (a) Influenza A virus (IAV) (accession numbers : MN220691-MN220698, data : SRR15305018 (33)) ; (b) Avian leukosis virus (ALV) (accession number : NC_001408, data : SRR7719537 (34)) ; (c) Japanese encephalitis virus (JEV) (accession number : AF045551.2, data : SRR11425577 (35)) ; (d) Human T-lymphotropic virus (accession number : NC_001436.1) ; (e) Hepatitis C virus (HCV) (accession number : AB047639, data : SRR27696424 (11)).

### RNA extraction and library preparation

RNAs were extracted from infected cells using TRI Reagent (AM9738 – Invitrogen), DNase I-treated as described in (16) and purified using RNA Clean and Concentrator (R1016 – Zymo Research). The library was then prepared by Novogene (Cambridge, UK), following their circRNA sequencing protocol. It includes 1) a circRNA enrichment by depletion of ribosomal and other linear RNAs through RNase R treatment; 2) a fragmentation of the circRNAs; 3) first-strand cDNA synthesis using random hexamers before the proper strand-specific library preparation.

### Data cleaning and bioinformatics analysis

To clean up the reads, we discarded the unmapped reads after mapping on the viral genome with the Burrows Wheeler Aligner (BWA) (36) under the command “bwa mem -a -T15 [fasta reference file] [fastq sequencing file]”. Next, all recovered RNA-seq data were cleaned using Trimmomatic (37) following the author’s protocol. It required trimming the adapters, eliminating very short reads (shorter than 70 nucleotides) and filtering bad quality reads (Sliding Window 4:15). The duplicated sequences resulting from PCR artifacts were removed using the package dedupe.sh (38). vCircTrappist is a pipeline relying on existing programs such as Samtools (39) and BWA followed by successive python scripts launched under a bash shell script. All the codes are available on the GitHub page of vCircTrappist (https://github.com/achasseu/vCircTrappist).

The first step (Figure 1A) of the pipeline is the alignment on the viral genome using BWA with the options “mem -a -T15”. The second step, run with splitfilter.py (Figure 1B), aims to isolate reads that exhibit splicing, removing those without at least one secondary alignment. The script analyzes flags in the alignment file, retaining reads that share the same ID, have one flag value at 2048 (or 2064 for reverse complement), and the other at 0 (or 16 for reverse complement). At this stage, reads mapping multiple times at various positions (flags 256 or 272 for reverse complement) are excluded, and the generated file is saved for subsequent user comparison.

The third step (Figure 1C), aiming at recovering the circRNAs, is carried out by several scripts, namely : circChaser.py, bsj_id.py, repet.py, circ_listing.py and circ_caracterisator.py. circChaser.py eliminates the reads originating from linear splicing. It analyzes the positions of reads with the same ID, identifying circRNA signatures. Specifically, if the start of a read is at position X, the end of the same read should be at position X-Y if it matches the sense strand, Y being an integer higher than 135 and smaller than 100,000. If it matches the antisense strand, the end of the read should be at position X+Y. This script produces a list of BSJ with their associated donor and acceptor backsplice sites (BSS).

bsj_id.py and repet.py, compare all BSJ and BSS to count backsplicing events. A backsplicing event is considered if two mapping reads have more than 95% identity conserved in their BSS sequences. During this analysis, donor and acceptor BSS of each event are compared to eliminate potential PCR or misalignment artifacts. Stretches of unique nucleotides or repetitive sequences like telomeric repeats, are removed at this step. Backsplicing events occurring more than once undergo a conclusive analysis with circ_listing.py and circ_caracterisator.py, characterizing them by identifying specific GT-AG motifs, which correspond to canonical splicing, at the BSS locations. The proximity of these sequences to the BSS attributes a score, which we utilize to categorize circRNAs as either canonical or non-canonical. For each BSS, a score of 0.5 is attributed if a “GU” (donor site, “AC” for reverse complement) or “AG” (acceptor site, “CU” for reverse complement) is located precisely at the splicing site. These sequences were chosen because they are conserved in the canonical splicing context. For each nucleotide between the splicing site and the nearest GU or AG sequence, the score is reduced of 0.1. Then the sum of the donor and acceptor scores (<= 1) is stored. Subsequently, this information is processed to generate four alignment files classifying backsplicing events based on their canonical status (score > 0.6) and further segregating them according to strand specificity.

These alignments are subsequently compiled using Samtools with the instructions “depth -a [FILENAME]” to generate a coverage table and Matplotlib to generate coverage plots and circRNA plots with covvisualisator.py and circ_visualisator.py (Figure 1D).

For the comparison with CIRI2, we followed the instructions provided by the publisher. This involved mapping the reads with BWA using the options “mem -a -T18” and utilizing both files of the paired-end sequencing. Subsequently, the resulting table was used to retrieve the circRNA reads from the alignment files. Samtools and a Matplotlib subscript (included in the vCircTrappist GitHub) were then employed to generate the coverage plots.

### RT-PCR confirmation

Reverse transcription was performed using the SuperScript IV kit (18090010 – Invitrogen) with random (250 nM; S1330S – New England Biolabs) or specific (100nM; Eurogentec) primers following the manufacturer’s recommendations. For each sample, a non-RT negative control was produced.

The PCRs were carried out using primers detailed in table 1 and the GoTaq G2 kit (M7841 – Promega) or the Q5 taq polymerase (M0491L – New England Biolabs). The cycles were adapted to the manufacturers’ instructions. All the positive PCRs were purified using the Nucleospin Gel and PCR Clean-Up kit (740609 – Macherey-Nagel), cloned into a pGEM-T easy vector using T4 DNA ligase (A1360 – Promega) using the TG1 strain of E.coli. Clones were sequenced by Sanger sequencing (Eurofins Genomics) for sequence confirmation.

**Table 1.**
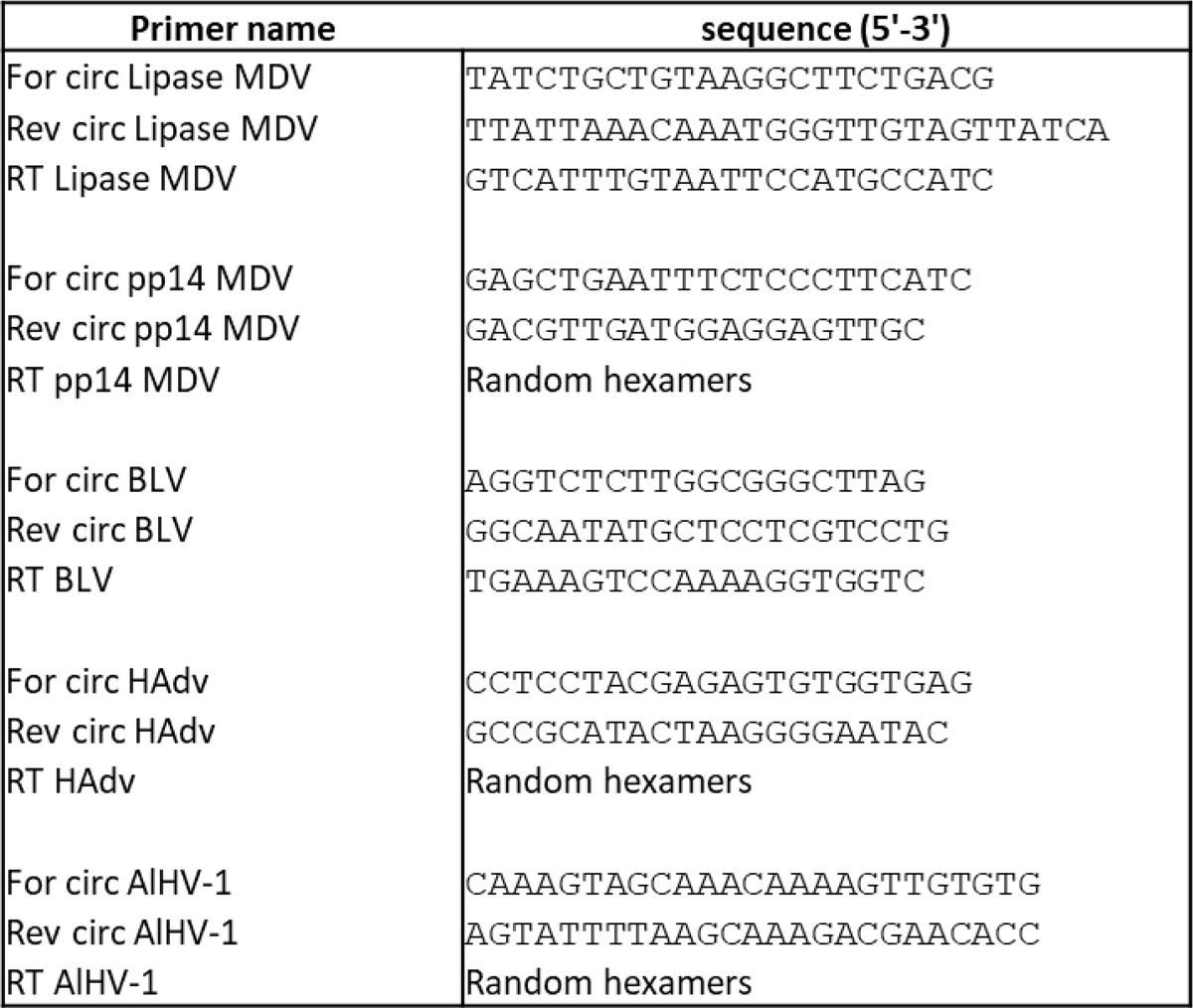
Primers used to confirm the vCircTrappist results.

### Grammatical and phrasing corrections

ChatGPT 3.5 (OpenAI) was used for grammatical purposes. No data, interpretation or texts were generated de novo using this tool.

**Figure 1.**
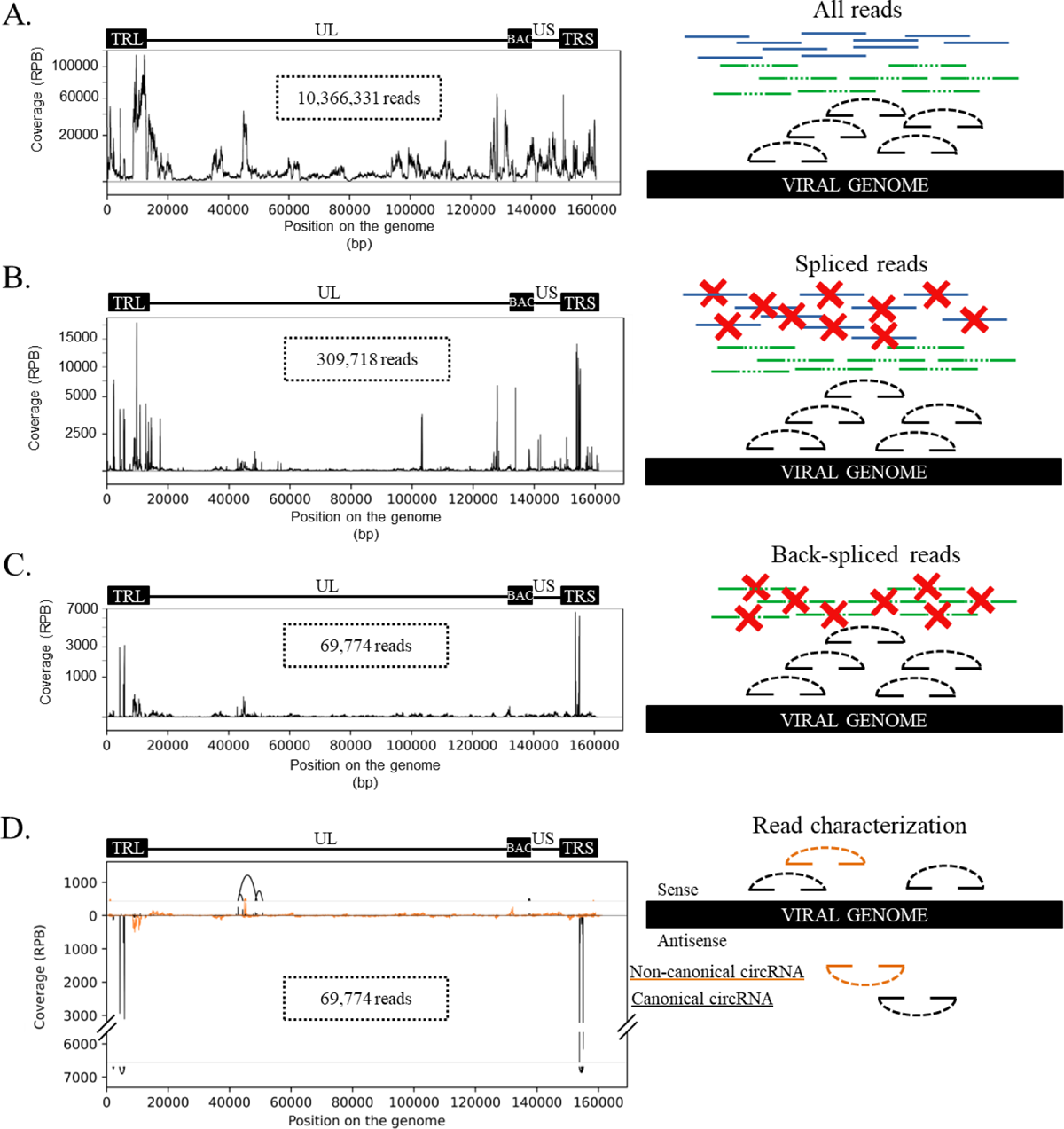
Description of the vCircTrappist process. (**A**) Mapping on the viral genome. The whole set of reads is mapped to the viral genome using BWA. (**B**) Sorting the spliced reads. The reads that present a splicing signature were recovered while the others were deleted. (**C**) Sorting the backsplice junction spanning reads. (**D**) Read characterization. The reads were sorted according to their features such as their splicing pattern or the strand they were originating from. The reads that correspond to a canonical U2 splicing pattern are depicted in black, while the non-canonical splicing reads are depicted in orange. The curves above and below the graph represent the 20 most abundant circRNA candidates found in the dataset. (**A-D**) On the left panels, the genome structure of the viral bacmid used for the transfection is depicted above the coverage plots. The genome is composed of two terminal repeats. The internal repeats were removed by the insertion of the bacmid cassette. The unique regions, short and long, are depicted. The X axis indicates the position on the viral genome (bp). The Y axis represent the normalized coverage. The normalization was made on the number of reads mapped at a specific locus divided by the total number of reads mapping on the viral genome, multiplied by one billion. On the right panels, the genome is depicted in black while the reads are depicted in green (linear splicing reads), blue (non-spliced reads) and black or orange (backsplice junction spanning reads).

## Results

### vCircTrappist: A Bioinformatics Pipeline for Viral circRNA Discovery

In our pursuit of discovering novel circRNAs, we have initially turned to widely employ circRNA identification programs. However, we encountered inconsistencies between the predictions of these programs and our experimental PCR results (see (16)). This prompted us to develop vCircTrappist, a dedicated tool designed to identify previously unrecognized circRNA candidates expressed by viruses during their infectious cycle.

The vCircTrappist pipeline hinges on the detection of backsplicing signals, as illustrated in Figure 1. The workflow entails several crucial steps:

1) Data Preprocessing: Prior to analysis, thorough preprocessing of raw sequencing data is essential. Data cleaning, including the removal of PCR duplicates, is performed after alignment to the viral genome with BWA (Figure 1A). This step is particularly critical in preventing the overestimation of circRNA quantities in subsequent analyses. Another essential part of the preprocessing is the manual preparation of the reference sequence. In Figure 1, working on an MDV infection, we opted to eliminate genomic duplications that could hinder proper read filtration. However, we retained short repeated regions. At this step, we obtained 10,366,331 reads mapping to the MDV genome after sequencing and analyzing an *in vitro* productive infection. 2) Spliced Read Filtering (Figure 1B): The initial stage of vCircTrappist involves the filtering of spliced reads, irrespective of their splicing patterns. This process entails identifying individual reads that map uniquely to a single genome or genome segment and exhibit at least one secondary alignment. Reads mapping on repetitive genome regions are discarded at this stage. They are filtered based on the fact that they comprise more than one primary alignment. After this filter, we recovered 309,718 reads from the 10 millions of the previous step. 3.1) Backsplicing Signature Detection: Subsequently, the precise mapping positions of primary and secondary alignments are leveraged to discern the backsplicing signature within the reads. Reads displaying a secondary alignment upstream in the viral genome sequence while being located downstream in the read sequence are considered to span a BSJ. These reads are saved, and their corresponding BSS and BSJ locations are recorded for further analysis. 3.2) Filters Application: Several filters are applied based on predefined criterias. Reads containing large repetitive elements such as stretches of unique nucleotides or telomeric repeats are removed. 3.3) Junction Counting: In the following process, BSJ and donor/acceptor BSS are compared between the extracted reads to obtain a raw count of identified junctions. 3.4) Candidate Selection (Figure 1C): Junctions that are represented more than once are used to extract reads more likely to be the results of genuine backsplicing. After all these filters, we recovered 69,774 circRNA reads. 4.1) Read Sorting (Figure 1D, table 2): The reads are sorted based on the strand from which they originate and their backsplicing pattern. Canonical U2 was assigned when the BSJ signature followed the conserved motif GT-AG. Non-canonical splicing was assigned when the BSJ did not follow this conserved signature. At this step, vCircTrappist generates a table (Table 2) which offers information to the users. They include the BSS and BSJ sequences ; the raw count of reads mapping the BSJ ; the sense and splicing pattern (canonical or not) of the circRNA ; the position of the splicing sites ; the expected size of the circRNA ; the gene locus from which originates the circRNA ; the longest common sequence between both splicing sites, to avoid false positive signals ; the segment on which the circRNA was mapped and ; the identifiers of the reads which encompass the BSJ. 4.2) Visualization (Figure 1D): A publication-ready graphical visualization is generated, depicting all the extracted reads.

Taken together, all those features make vCircTrappist a robust and tailored pipeline, purpose-built for the identification of viral circRNAs within the unique context of viral infections. Its adaptability to key viral features, including repetitive elements, transcriptional density of viral genomes, and the non-canonical patterns of circRNA expression, positions this pipeline as a potentially indispensable tool for the identification of viral circRNAs.

**Table 2.**
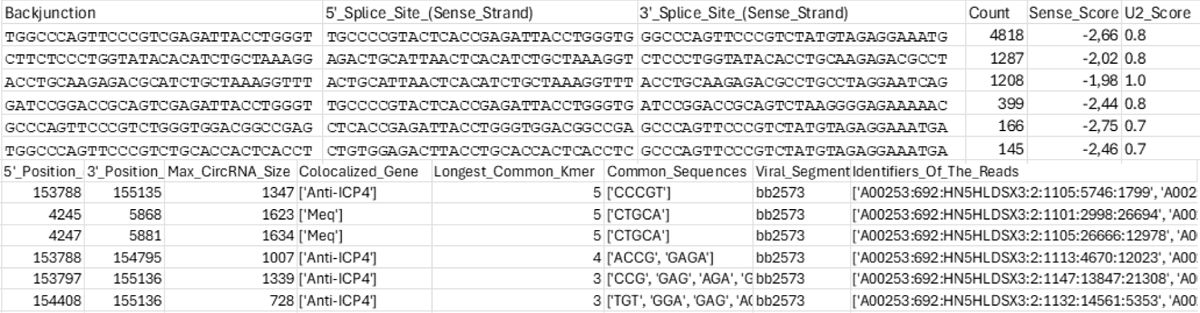
Example of dataset extracted using vCircTrappist. This example displays the data obtained for MDV, as in Figure 1, 2 and S1. The three first columns represent the backsplice junction and the backsplice sites. The « Count » column represent the raw number of reads mapping the backsplice junction without normalization. The Sense_score column is calculated based on the strand which is covered by the reads mapping the backsplice junction. The U2_score is calculated based on the distance of the backsplicing site from a canonical splicing site. The 5’ and 3’ position are the positions of the splicing sites and they determine the Max_CircRNA_Size. The « Colocalized Gene » column requires a completed GFF file to determine the colocalizing genes. The « Longest Common Kmer » was identified comparing the backsplice sites and the « Common Sequences » were extracted accordingly. The « Segment » is the viral segment on which the backsplice junction was identified. The identifiers of the reads are available in the SAM alignment file.

### vCircTrappist finds more circRNA candidates than CIRI2 from viral infections datasets

To validate the efficacy of vCircTrappist in viral circRNA identification, we conducted a comparative analysis using a newly acquired dataset from infected cells, juxtaposing its performance with the widely employed program, CIRI2 (19) (Figures 2 and S1). It is noteworthy that visualization of the CIRI2 results required the development of a Matplotlib subscript to provide graphical representation of BSJ reads. This subscript is available on the vCircTrappist GitHub page.

In our initial analysis, we identified a substantial number of viral circRNAs using both CIRI2 and vCircTrappist. To compare their performance, we chose to focus on the adenovirus infection model which we considered relevant for the comparison because we found abundant circRNAs with both vCircTrappist and CIRI2 (Figure 2A). By analyzing the coverage plots obtained from both pipelines, we observed numerous peaks detected by vCircTrappist but not by CIRI2. However, one circRNA appeared more abundant in the CIRI2 coverage plot than in vCircTrappist, although both programs detected it. Further investigation revealed a canonical splicing spot, highlighting the ability of both CIRI2 and vCircTrappist to recognize canonical circRNAs. We attribute the difference in abundance to vCircTrappist’s use of only one mate of the paired-end sequencing, while CIRI2 analyzes both mates. Our pipeline was designed without considering paired-end sequencing due to the RT-induced rolling-circle amplification of circRNAs, leading to concatemerization of circRNAs in the resulting cDNAs, as described in (11). This amplification may bias subsequent analyses when using paired-end sequencing, as both ends of the reads might map to the same BSJ.

To standardize the analysis for diverse infection contexts, we generated a heatmap by dividing into 20 fragments the viral genomes of five viral pathogens and comparing the mean coverage obtained by CIRI2 and vCircTrappist in the corresponding 100 fragments (Figure 2B). This approach allowed us to evaluate vCircTrappist’s efficacy compared to CIRI2. Notably, vCircTrappist identified more circRNA hits than CIRI2 in most cases, with 93 out of 100 genome fragments analyzed showing a higher number of circRNAs detected by vCircTrappist. Interestingly, CIRI2 identified more hits in a specific region in the adenoviral infection case (described above).

We then focused on a MDV fragment for which vCircTrappist detected abundant circRNA candidates generated from non-canonical splicing (Figure 2C). We designed inverted PCR oligos for two loci involved in viral pathogenesis and obtained RT-PCR amplicons in both cases (Figure 2D), confirming the relevance of the signals obtained with vCircTrappist.

Collectively, these results highlight the vast wealth of information provided by vCircTrappist when compared to CIRI2, the current bioinformatics pipeline most frequently used for the identification of viral circRNAs. Notably, the circRNAs newly identified using vCircTrappist are linked to genes implicated in viral pathogenesis, further emphasizing vCircTrappist’s potential to shed light on crucial circular transcripts.

**Figure 2.**
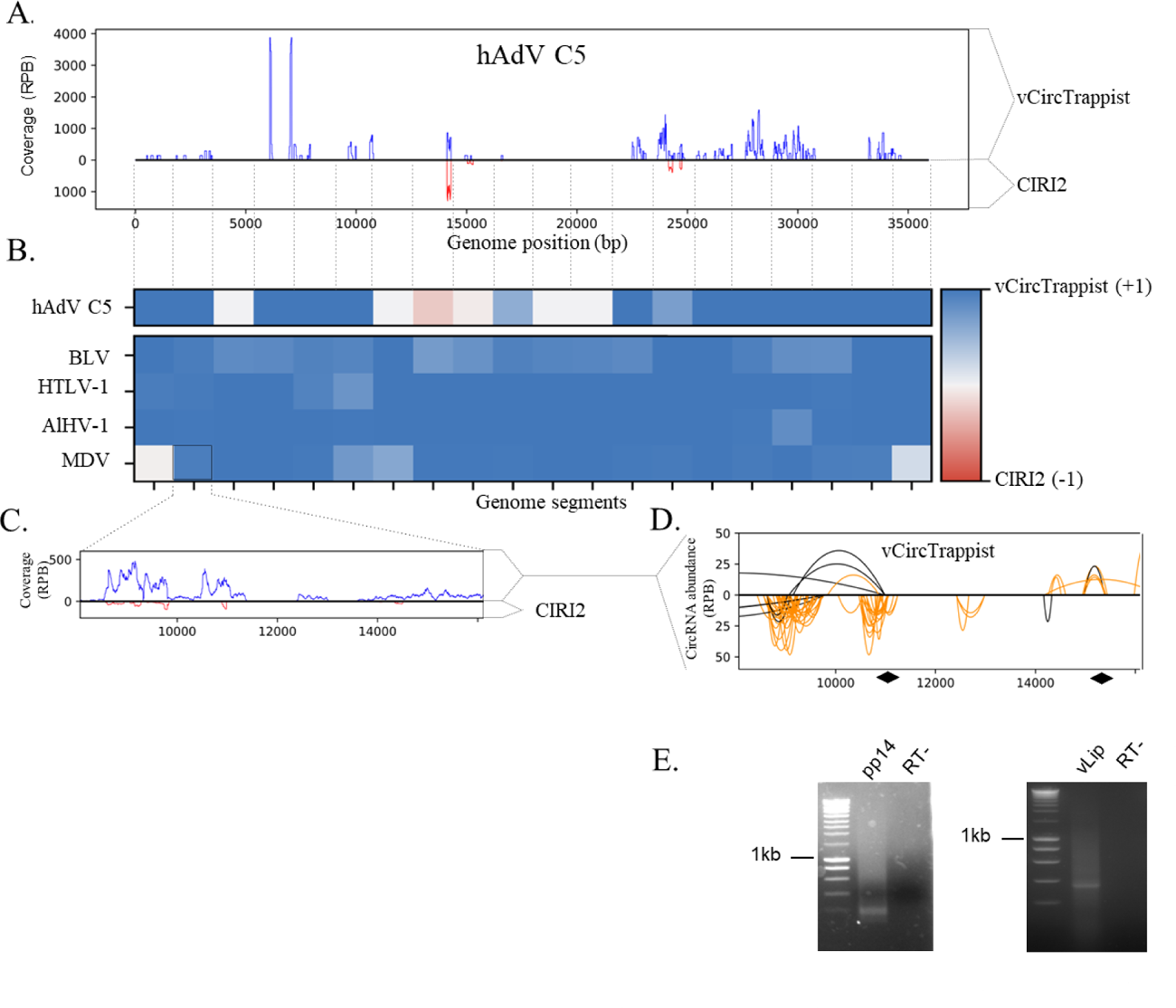
Comparison of vCircTrappist with CIRI2. (**A**) vCircTrappist vs CIRI2 coverage of circRNA reads on the hAdV C5 genome. vCircTrappist reads are mapped to the upper part of the graph while CIRI2 reads are mapped below. The Y axis represents the coverage in circRNA reads per billion reads mapped to the viral genome. The X axis represent the positions on the genome in base pairs. (**B**) Comparison of vCircTrappist vs CIRI2 in multiple infection contexts. The heatmap represents the relative abundance of circRNAs on diverse viral genome segments. The viral genomes were segmented for graphical scale purposes. The segments are each 1/20 of the viral genome. The data were obtained as followed : (vCircTrappist coverage – CIRI2 coverage)/(vCircTrappist coverage + CIRI2 coverage). The results are on a scale from 1 (blue – vCircTrappist has a better coverage in this particular genome part) to -1 (red – CIRI2 has a better coverage). (**C**) Local mapping to ascertain the better coverage of vCircTrappist vs CIRI2 using the MDV model. (**D**) Analysis of the coverage obtained with vCircTrappist. The Y axis indicates the amount of time individual reads have mapped to a specific backsplice junction divided by the total number of reads mapping on the viral genome and then multiplied by one billion. The X axis indicates the positions (in bp) where the circRNA are backspliced. The position of the primers for subsequent RT-PCR confirmation are indicated in black. (**E**) PCR confirmation of the loci that were identified only by vCircTrappist. The smears show the abundance of alternative transcripts when it comes to circular RNAs. The ladder is the SmartLadder from Eurogentec, the 1kb band was indicated on the Figure.

### vCircTrappist Discovers Novel Viral circRNAs

Having established the competence of vCircTrappist for the discovery of novel viral circRNAs, we proceeded to apply it to both new (Figures 3; S2-S4) and existing datasets (Figures S5-6). In this comprehensive investigation, we delved into the realm of circRNAs expressed by a wide spectrum of DNA and RNA viruses.

We applied vCircTrappist to three infection contexts. On the one hand, we explored circRNA profiles in two animal models associated with virus-induced lymphoproliferation: lymphocytes collected and amplified from infected animals with alcelaphine herpesvirus 1 (AlHV-1) and bovine leukemia virus (BLV) (Figures 3A–F; S2). On the second hand, we applied the pipeline in the context of an adenoviral *in vitro* infection (Figures 3G-I ; S4), employing the human adeno virus C5 (hAdV C5), commonly used as a benchmark in human adenovirus studies. RNA extraction was carried out at the 18-hour post-infection time point. In these three viral infection contexts, vCircTrappist effectively identified a multitude of new circRNA candidates. To validate these findings, we confirmed one circRNA exhibiting high read coverage for each virus via inverse RT-PCR followed by Sanger sequencing. This approach, based on divergent primers exclusively amplifying circRNAs, added robustness to our circRNA identifications (Figures 3B ; 3E ; 3H).

The AlHV-1 infection revealed abundant circRNA expression through vCircTrappist analysis. We unveiled numerous circRNAs loci and one of them was subsequently confirmed through RT-PCR and Sanger sequencing (Figures 3B-C). One striking discovery was a locus (labeled “a” on Figure 3A) previously unassociated with any known transcripts or open reading frames (ORFs). While the circRNAs associated with this locus did not adhere to a canonical U2 splicing pattern, the peak signature in the coverage plot revealed conserved exonic profile observed from two independent *in vivo* samples. This suggests that this highly expressed locus might be the source of non-coding circRNAs possibly involved in virus-induced pathogenesis. A second locus (labeled “b” on Figure 3A) is linked to the expression of the ORF73 protein, responsible for maintaining the episomal viral genome within the infected cell (31, 40). Collectively, the results obtained from AlHV-1 infection highlighted the production of circRNAs of viral origin in another viral induced lymphoproliferative disease.

In the context of a retroviral infection; similar results were obtained by analysing the viral circRNAome. Three different models of retroviral infection by BLV were used: two of these were tumor-derived BLV-infected cell lines constitutively producing high levels of viral transcripts (YR2_LTaxSN_ and L267_LTaxSN_) while the third one was an in vitro model of BLV infection generated by transfection of human-derived uninfected 293T cells with a plasmid expressing a complete and infectious BLV clone. The obtained data presented intriguing features as they lacked a specific splicing pattern and conserved splicing sites (Figure 3D). However, they exhibited exceptional reproducibility in three distinct models of productive viral infection (Figures 3D ; S2). Data obtained before reactivation did not reveal any circRNA expression during latency, most probably due to a reduced level of viral transcripts (data not shown). Nevertheless, the consistency across the three sequencing results from productive infections established a key point: the generation of circRNAs was not contingent on the cell type. The major circRNA expression peak was situated at the pol gene, which encodes for the viral reverse transcriptase. Utilizing divergent PCR, we confirmed the presence of circRNAs originating from this region in the two most relevant infection models (L267_LTaxSN_ and YR2_LTaxSN_) (Figures 3E-3F). We also carried out a circRNA sequencing on Human T Lymphotropic Virus 1 (HTLV-1)-infected cell lines (Figure S3) since BLV stands as a robust animal model for this human retrovirus. This revealed strong coverage peaks in the 3’ end of the HTLV-1 genome in the vicinity of the tax and rex genes.

In the context of the infection with hAdV C5, we identified three loci (Figure 3G) from which circRNAs were abundantly detected. These loci exhibited distinctive features: 1) the first locus (labeled “a” on Figure 3G) appeared to be a stabilized intron between the TPL1 (Tripartite Leader 1) and TPL2 (from position 6090 to 7085), as its backsplicing pattern exhibited the typical signature of introns; 2) the second locus (labeled “b” on Figure 3G) consists on a canonical exonic circRNA made from the third exon of E2B transcript (from position 14299 to position 14111); 3) the last locus (labeled “c” on Figure 3G) produced multiple non-canonical circRNAs that mapped to the forward strand of the E3 genes. The circRNA produced from locus “b” was validated by inverse RT-PCR (Figure 3H).

These findings collectively revealed, for the first time, the expression of circRNAs produced during an adenoviral infection of human cells. By examining multiple time points (12 hours, 18 hours, and 24 hours post-infection; Figures 3H and S4), we also characterized the expression of viral circRNAs throughout the course of the infection. As expected, circRNA expression mirrored the dynamic progression observed in linear RNAs. For example, the stabilized intron mapped between the TLP1 and 2 (locus “a”) exhibited a remarkable increase in expression concomitantly with the expression of the late transcripts (Figure S4, time points 18 and 24 hours post infection).

In these infection contexts, as well as for the BLV infections, we were able to obtain multiple PCR amplicons. These multiple bands on the electrophoresis results correspond to the RT-induced rolling circle amplification of circRNAs as discussed in the previous paragraph. More precisely, the circRNA of the locus “b” in Figure 3G is amplified with a primer pair that should lead to a band located at a size of 157bp in Figure 3H if we do not consider the RT-induced rolling circle amplification. Considering it, every upper bands correspond to 157bp added to multiple times the size of the circRNA, 187bp. We therefore obtain bands at 157bp, 344bp, 531bp, 718bp and 905bp all corresponding to the same circRNA. This is again a strong proof-of-concept for vCircTrappist to be able to identify accurately circRNAs.

Collectively, these results affirm vCircTrappist as an exemplary tool for the identification of viral circRNAs within diverse infectious contexts. Importantly, we successfully unveiled new circRNAs in all studied viruses. In the context of this study, we also investigated four viral circRNAome using preexisting datasets (Figures S5-S6). In all cases we identified numerous circRNA candidates. More precisely, we were able to identify 1023 unique BSJs for the Influenza A virus (IAV) (Figure S5, Table 3), while we obtained 135 for the Avian Leukosis virus (ALV) (Figure S6A, Table 2), 2764 for Japanese Encephalitis Virus (JEV) (Figure S6B, Table 3) and 208 for HCV (Figure S6C, Table 3). Writing these lines, we have no doubt that vCircTrappist will emerge as the gold standard in viral circRNA identification.

**Figure 3.**
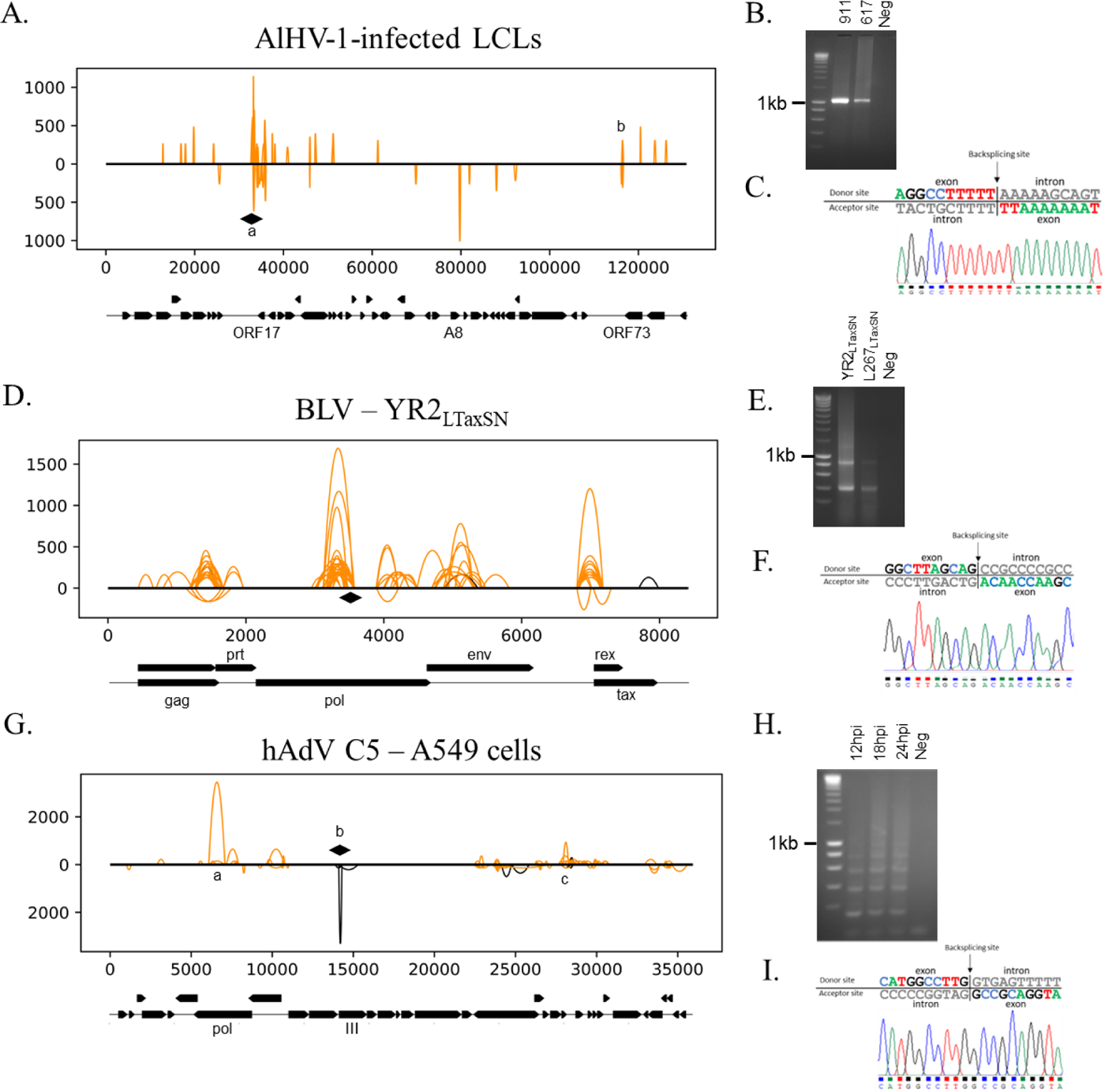
Investigation of unexplored viral circRNAs from diverse infections. (**A**) Viral circRNAs expressed from an *in vivo* infection with AlHV-1. A lymphoblastoid cell line (LCL) from *in vivo*-infected calves was recovered and grown in culture. (**B**) PCR confirmation of the results obtained by Illumina Sequencing. The lanes represent two different infected animals and a negative control. The bands were purified and cloned into a vector to be sequenced thanks to Sanger sequencing. (**D**) Viral circRNAs expressed from BLV productively-infected cell line YR2_LTaxSN_. (**E**) PCR confirmation of the results obtained by Illumina Sequencing. The lanes represent two different cell lines obtained from *in vivo* infected sheep. (**G**) Viral circRNAs expressed from an *in vitro* infection with hAdV-C5. After infection, RNAs were extracted after 18h post infection. (**H**) PCR confirmation of the results obtained by Illumina Sequencing. The lanes represent three different time-points of infection. (**A,D,G**) The 100 more expressed circRNAs are depicted on the graphs. The X axis represent the position on the viral genome. The Y axis indicates the amount of time individual reads have mapped to a specific BSJ divided by the total number of reads mapping on the viral genome and then multiplied by one billion. The black lines represent the canonical circRNAs. The orange lines represent the non-canonical circRNAs. The height of the curves represent the normalized abundance of each individual backsplice junction. The lines up from the genome are mapped in the same sense than the viral genome while the lines under are antisense to the viral genome. The relevant ORFs of the viral genome are depicted under the graphs. For graphical purposes, the splicings of the tax and rex genes were not depicted. The loci targeted for RT-PCR confirmation are indicated by black losanges. Peak of interest described in the text are indicated by lowercase letters. (**B,E,H**) The ladder is the SmartLadder from Eurogentec. The 1kb band was indicated next to the gel picture. (**C,F,I**) Chromatogram obtained from Sanger sequencing of the corresponding candidate circular RNA.

**Table 3.** List of viruses for which data was obtained using vCircTrappist. The samples marked by an asterisk were acquired in the context of the present study.

| Virus | Number of unique backsplice junctions |
| --- | --- |
| Marek's disease virus (MDV)* | 8453 |
| Alcelaphine Herpesvirus 1 (AIHV-1) - Cow 1* | 302 |
| Alcelaphine Herpesvirus 1 (AIHV-1) - Cow 2* | 3410 |
| Human Adenovirus C5 (hAdV C5) - 12h p.i.* | 13 |
| Human Adenovirus C5 (hAdV C5) - 18h p.i.* | 87 |
| Human Adenovirus C5 (hAdV C5) - 24h p.i.* | 404 |
| Influenza A virus (IAV) – SRR15305018 | 1023 |
| Avian Leukosis Virus (ALV) – SRR7719537 | 135 |
| Japanese Encephalitis Virus (JEV) – SRR11425577 | 2764 |
| Bovine Leukemia Virus (BLV) - YR2<sub>LTaxSN</sub>* | 882 |
| Bovine Leukemia Virus (BLV) – transfected 293T* | 752 |
| Bovine Leukemia Virus (BLV) – L267<sub>LTaxSN</sub>* | 468 |
| Human T Lymphotropic Virus 1 (HTLV-1) – HUT102* | 147 |
| Human T Lymphotropic Virus 1 (HTLV-1) – SLB1* | 427 |
| Hepatitis C Virus (HCV) - SRR27696424 | 208 |

## Discussion

In this study, we have introduced vCircTrappist, a novel bioinformatics pipeline meticulously designed for the comprehensive exploration of circularization events within viral transcripts. By imposing rigorous filters, we successfully isolated these distinct sequences from both new and existing datasets. When compared to CIRI2, the current gold standard for circRNA identification, vCircTrappist demonstrated superior accuracy and robustness in detecting circRNA candidates with potential implications for the pathogenesis process. Therefore, these loci deserve further characterization. Although other programs were assessed (20, 21), they were omitted from our analysis as they exhibited similar limitations to CIRI2 in the context of viral circRNAs. Notably, the fact that 1) they are based on paired-end sequencing that tend to overestimate the abundance of circRNAs and 2) they all rely on the recognition of canonical splicing patterns. Our findings, particularly from adenoviral, herpesviral and retroviral infections, are of significant note, underlining the potential of vCircTrappist to become the future gold standard for viral circRNA identification.

Our results offer promise in the development of an open, unbiased, and biologically relevant approach for viral circRNA discovery. While the concept of identifying BSJ-covering reads is not new (19–21), the recent characterization of numerous non-canonical circRNAs (10, 12–14, 16) necessitates a departure from the stringent parameters imposed by current programs. In our comparative analysis, vCircTrappist consistently found more hits than CIRI2 across various viral infection contexts. While CIRI2 is acknowledged as the gold standard for cellular circRNAs, only vCircTrappist demonstrated precise mapping of numerous non-canonical circRNAs on viral genomes. This enhanced detection can be attributed to our choice of employing diverse criteria for circRNA identification, applying custom filters tailored to the unique attributes of viral genomes. For example, we deliberately excluded reads spanning repeated regions to eliminate potential inaccuracies. This is because many viruses harbor large repeated sequences in their genome for regulatory purposes. In this context, we can cite the Long Terminal Repeats (LTR) in retroviruses or the diverse repeat regions in herpesviruses. These custom filters, although increasing computational time, significantly enhance the generation of publication-ready data, as illustrated in Figure 2. However, it is crucial to note that these repeated regions may serve as sources of circRNAs, and with current technologies, distinguishing them from linear RNAs is challenging. In our case, to study genes located in the inverted repeats of MDV, we kept only one of these copies and suggest the reader to proceed in the same manner.

Our newfound data concerning three distinct viruses (AlHV-1; BLV; hAdV C5) provides valuable insights into the strengths and potential limitations of vCircTrappist thanks to numerous RT-PCR confirmations of circRNA candidates. During our investigation of hAdV C5 circRNAs, we encountered a circRNA candidate that, upon further analysis, was determined to be an intron-derived transcript. Yet, its expression pattern suggested that the transcript was stably expressed throughout the infection course, which, in itself, was intriguing and worthy of further investigation. This finding illustrated that vCircTrappist could identify transcripts beyond traditional circRNAs, highlighting its potential for serendipitous discoveries.

In both BLV and hAdV C5 data, we observed circRNAs encoded in the vicinity of the pol genes. While these two genes use different mechanisms to generate new viral (pro)genomes, these observations might reflect the role of circRNAs in genome maintenance and/or duplication. We hypothesize that the circRNAs produced from these spots might interact with the proteins produced from the same genes and alter the processing of the nascent (pro)genome. However, we were not able to confirm this hypothesis, and more work will be required in this context.

The case of AlHV-1 also warrants attention. While the identification of abundant non-coding transcripts during a herpesviral infection is not unexpected (16, 41, 42), the primary expression peak is positioned at a locus that has not been previously characterized, as it is not associated with any ORFs. However, it is in close proximity to the expression locus of the viral microRNAs (miRNAs). Although this locus was not linked to the pathogenesis of AlHV-1 (32), it may act concomitantly with the newly described circRNAs or, conversely, mitigate their effect. Previous research on the Epstein-Barr Virus (EBV) (9) has shown that the circRNA associated with the Latent Membrane Protein 2A sponges a miRNA involved in p53 regulation, a regulatory cycle linked to the progression of gastric cancer. Another possibility is the involvement of these circRNAs as non-coding transcripts essential for the establishment of latency, as demonstrated for diverse alphaherpesviruses (43–45). In summary, the discovery of circRNAs produced during AlHV-1-induced lymphoproliferation may offer a fresh perspective on pathogenesis.

Our extensive analysis of diverse viruses highlights the potential impact of vCircTrappist on the virology community. Along with previous studies (10, 12–14, 16, 17, 22), we once again observe that viral circRNAs are primarily produced in non-canonical ways. While we cannot provide a definitive mechanistic explanation for this phenomenon, previous research (17) has identified the involvement of the exonuclease activity of the NSP14 protein in the generation of these transcripts during a coronavirus infection. It is worth noting that this mechanism might be more universal and could involve both viral and cellular factors, as non-canonical circRNAs have also been identified in plant and metazoan non-infected cells (46, 47).

In summary, we present here a pioneering bioinformatics pipeline with the potential to unveil previously uncharted roles within viral transcripts characterized by diverse splicing patterns. Altogether these analyses allowed us to detect an important number of unique BSJ from various viral infection models as presented in Table 3. The study opens the door to an intricate field that needs thorough exploration. It began with the description of new circRNAs from *in vitro* and *in vivo* infections (Figures 2 and 3). Then we analyzed their production during an infection kinetics (Figure S4). And finally, we explored existing datasets with the ALV, HCV and JEV (Figure S6) and with the different segments of the IAV (Figure S5). Nonetheless, the circRNA landscape of the vast majority of viruses remains to be explored.

## Data availability statement

vCircTrappist will be released after publication of this paper (https://github.com/achasseu/vCircTrappist). All data are available upon request to address to D.C. or will be available on a GEO repository after publication of this paper.

## Acknowledgments

We gratefully thank Laetitia Wiggers (UNamur) for her help with the PCR assays. This study was funded by the Belgian National Fund for Scientific Research (FRS-FNRS, Belgium) through its Projets de Recherche (PDR) program awarded to BM, BD and CVL [grant number T.0160.23]. Three Fonds pour la Recherche dans l’Industrie et l’Agriculture (FRIA) scholarships were awarded to AC [grant number 40014728], MB [grant number 35484055] and SM [grant number 40021558], respectively. A Chargé de Recherche fellowship was awarded to CY [grant number FC5555]. A EU Marie Sklodowska-Curie COFUND Action was awarded to MG [Grant number 801505].

## Supplemental Figures

**Figure S1.**
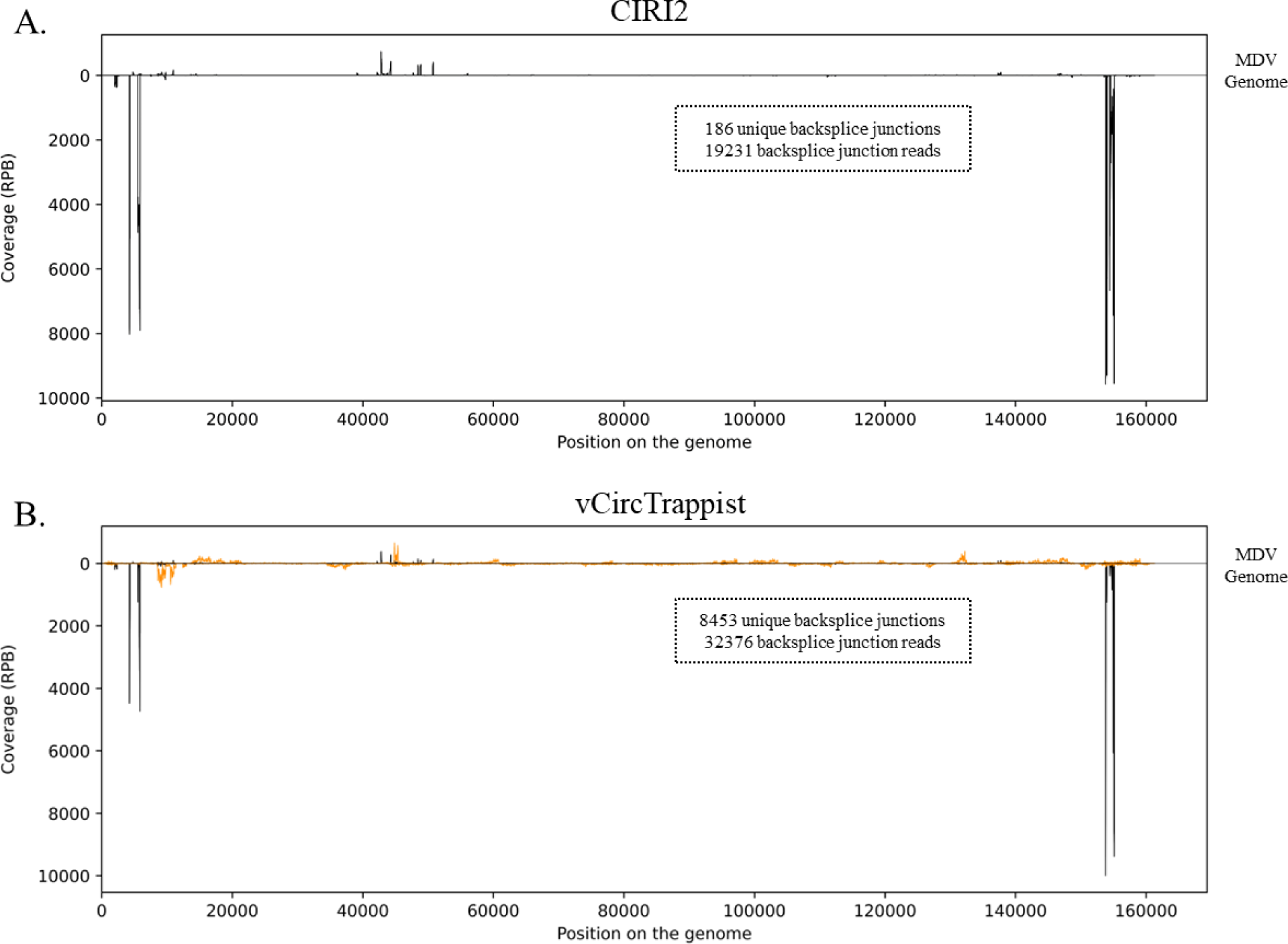
vCircTrappist comparison with CIRI2. (**A**) CIRI2 mapping of circRNAs on the MDV genome. In that case, both reads of the pair were used. (**B**) vCircTrappist mapping of circRNAs on the MDV genome. The orange peaks represent non-canonical backsplice junction while the black peaks represent a canonical backsplicing pattern. The peaks represent the normalized coverage (backsplice junction spanning reads per billion mapped reads on the viral genome (RPB)).

**Figure S2.**
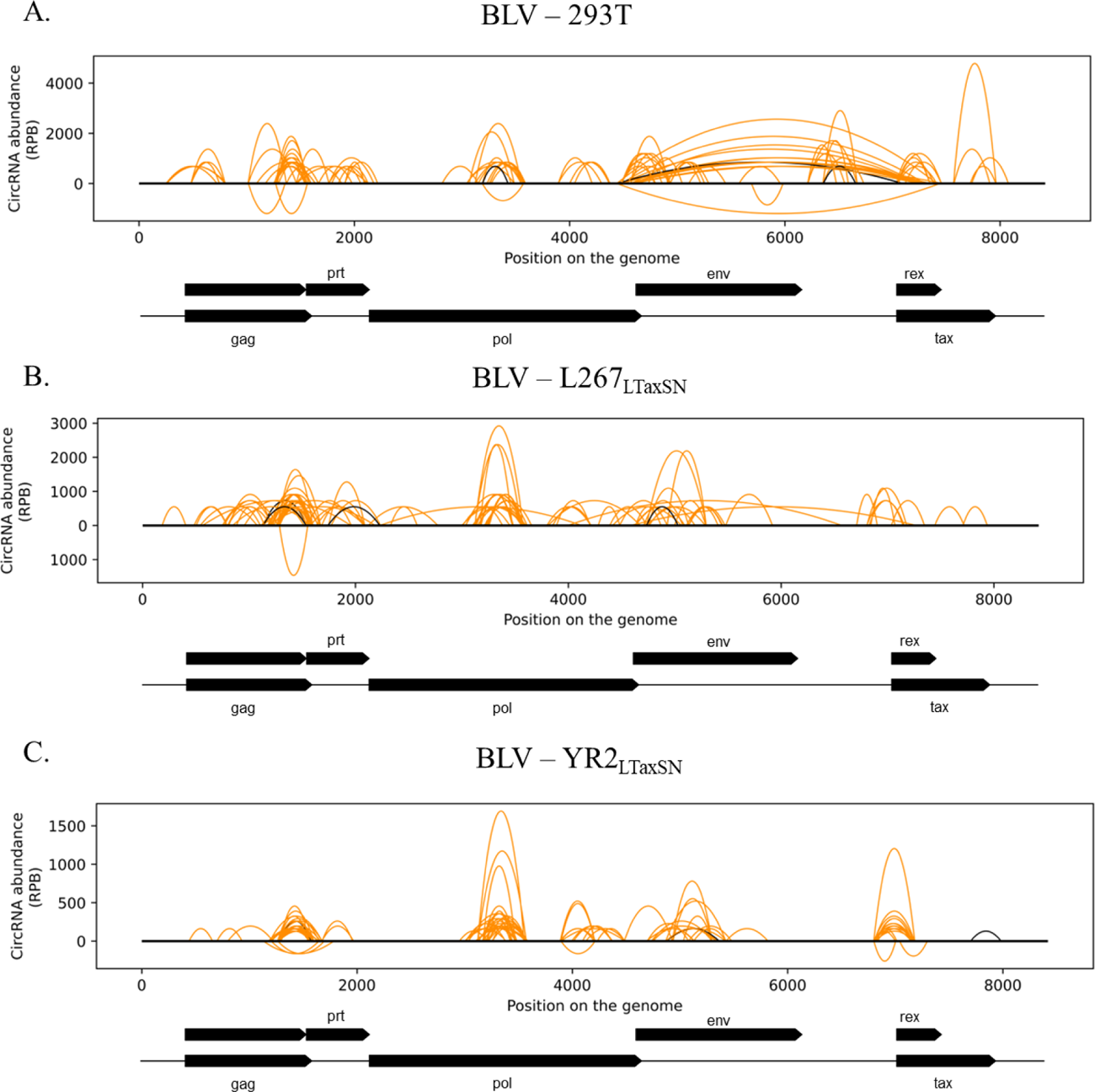
Viral circRNA idenfication in three models of BLV infections. (**A**) CircRNAs mapped to the BLV genome after a transfection of the viral genome in HEK 293T cells. (**B**) CircRNAs mapped to the BLV genome during a productive infection in the L267_LTaxSN_ cell line. (**C**) CircRNAs mapped to the BLV genome during a productive infection in the YR2_LTaxSN_ cell line. Relevant ORFs were indicated under the graphs on the same scale as the viral genome. For graphical purposes, the splicings of the tax and rex genes were not depicted. The Y axis indicate the abundance of unique circRNAs in reads mapping the backsplice junctions per billion of reads mapping on the viral genome.

**Figure S3.**
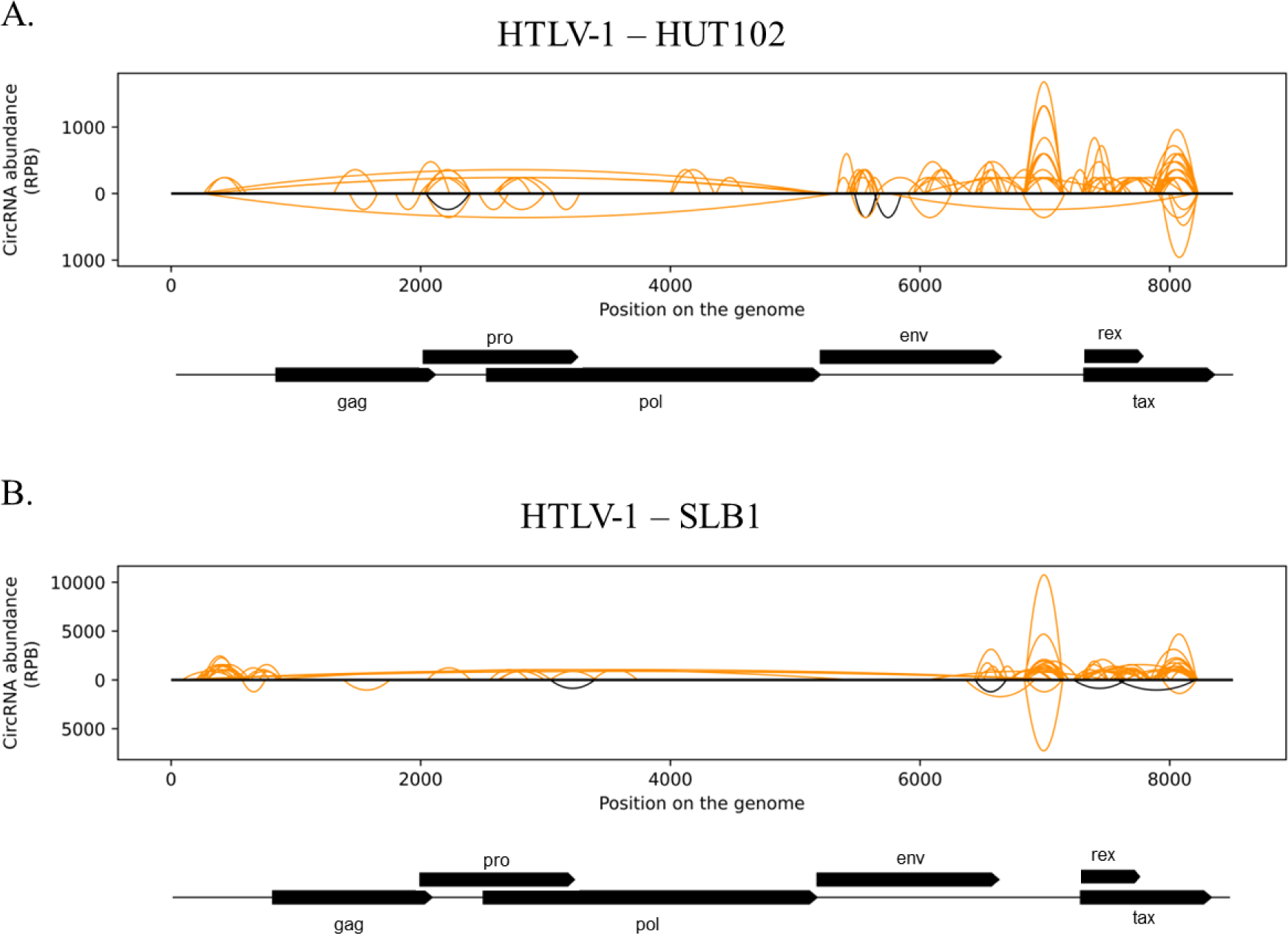
Viral circRNA identification in two HTLV-1 infection models. (**A**) CircRNAs mapped to the HTLV-1 genome of the productively-infected cell line HUT102. (**B**) CircRNAs mapped to the HTLV-1 genome of the productively-infected cell line SLB1. Relevant ORFs were indicated under the graphs on the same scale as the viral genome. For graphical purposes, the splicings of the tax and rex genes were not depicted. The Y axis indicate the abundance of unique circRNAs in reads mapping the backsplice junctions per billion of reads mapping on the viral genome.

**Figure S4.**
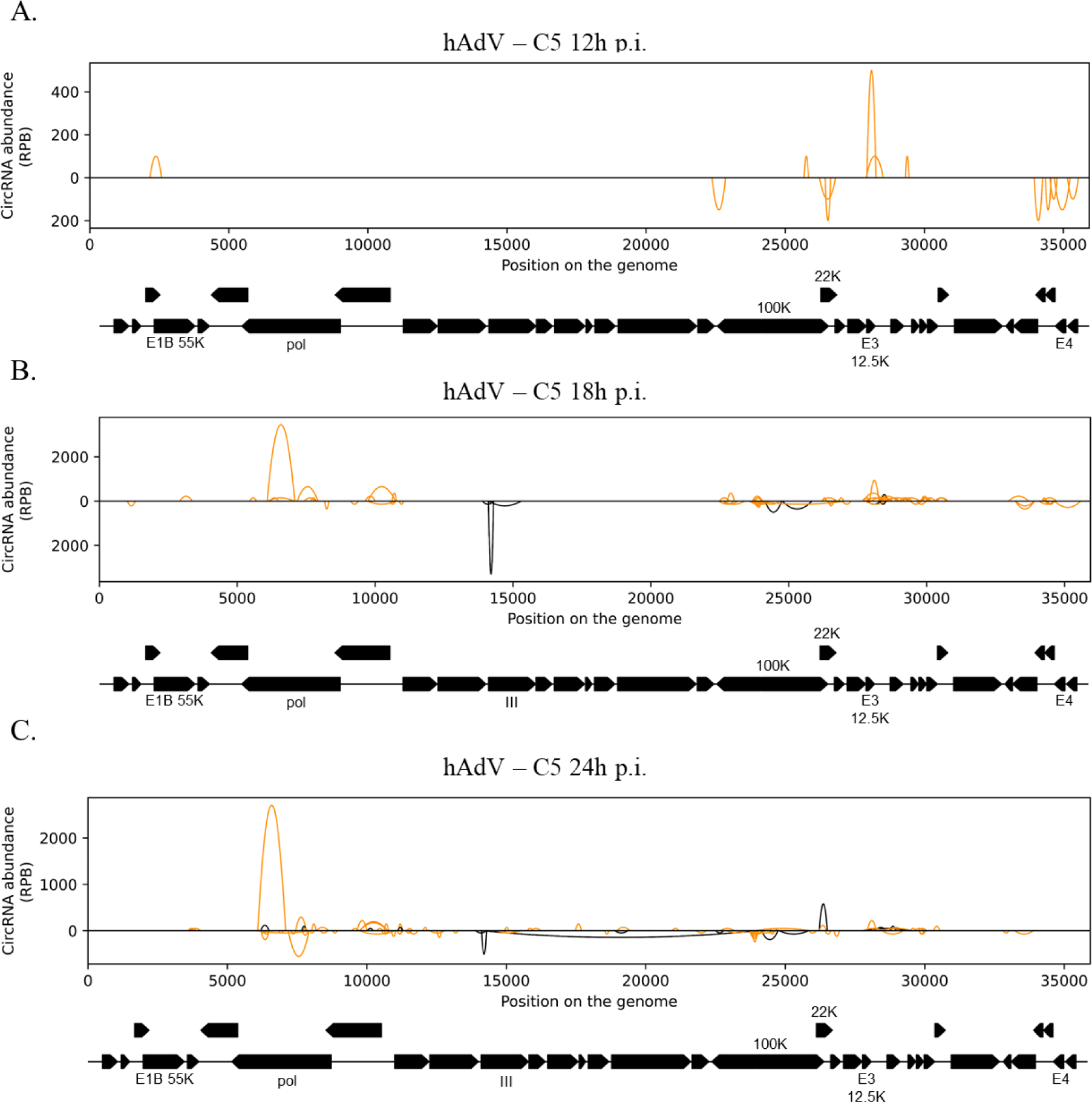
Viral circRNA identification during the course of an adenoviral infection. The circRNAs were mapped to the hAdV-C5 genome after 12h (**A**), 18h (**B**) or 24h (**C**) of infection. The infection was made with A549 cells. Relevant ORFs were indicated under the graph on the same scale as the viral genome. The Y axis indicate the abundance of unique circRNAs in reads mapping the backsplice junctions per billion of reads mapping on the viral genome.

**Figure S5.**
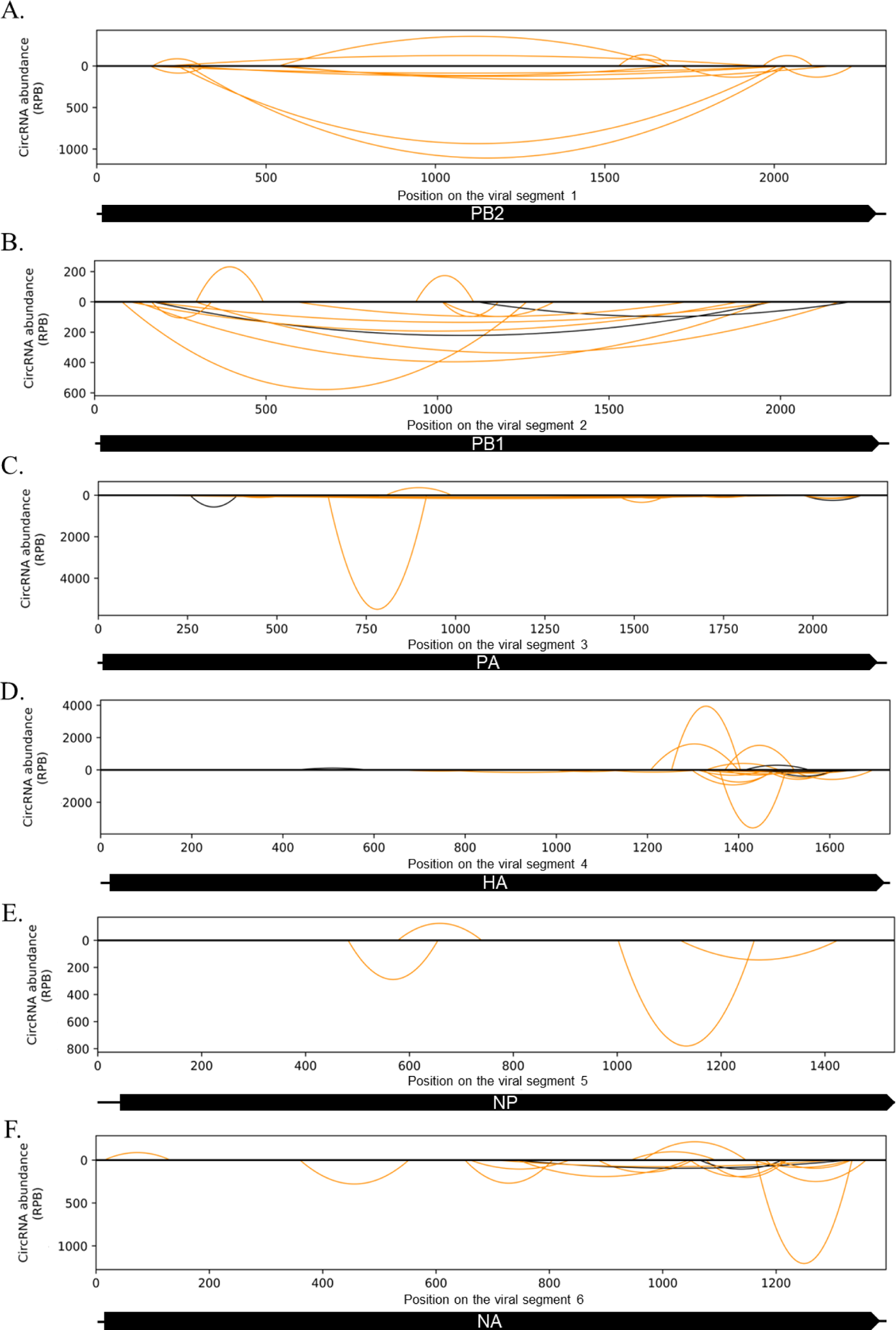
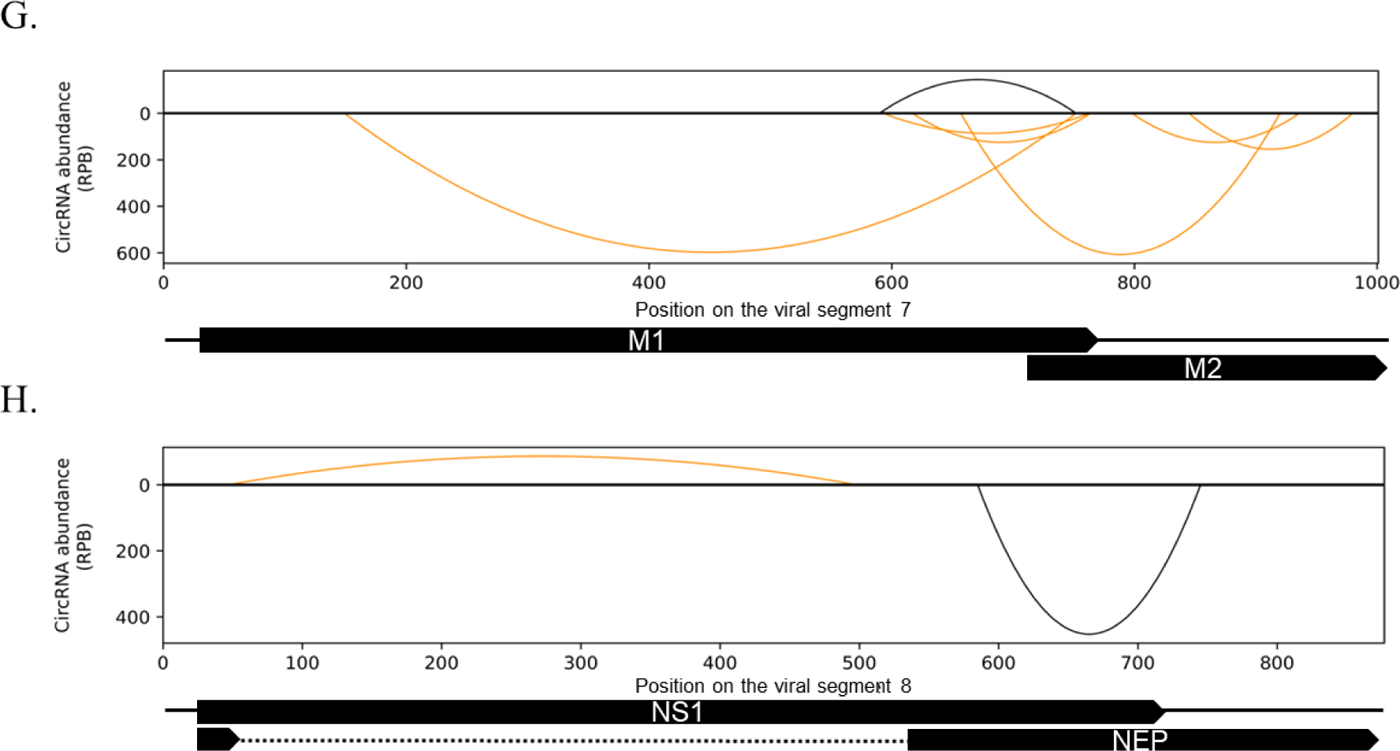
Viral circRNA identification from an IAV infection. (**A-H**) A549 cells were infected with the H1N1 strain of IAV in the study of Min *et al*, 2023. The eight segments of the virus were processed through vCircTrappist and depicted in the Figure. The Y axis indicate the abundance of unique circRNAs in reads mapping the backsplice junctions per billion of reads mapping on the viral genome.

**Figure S6.**
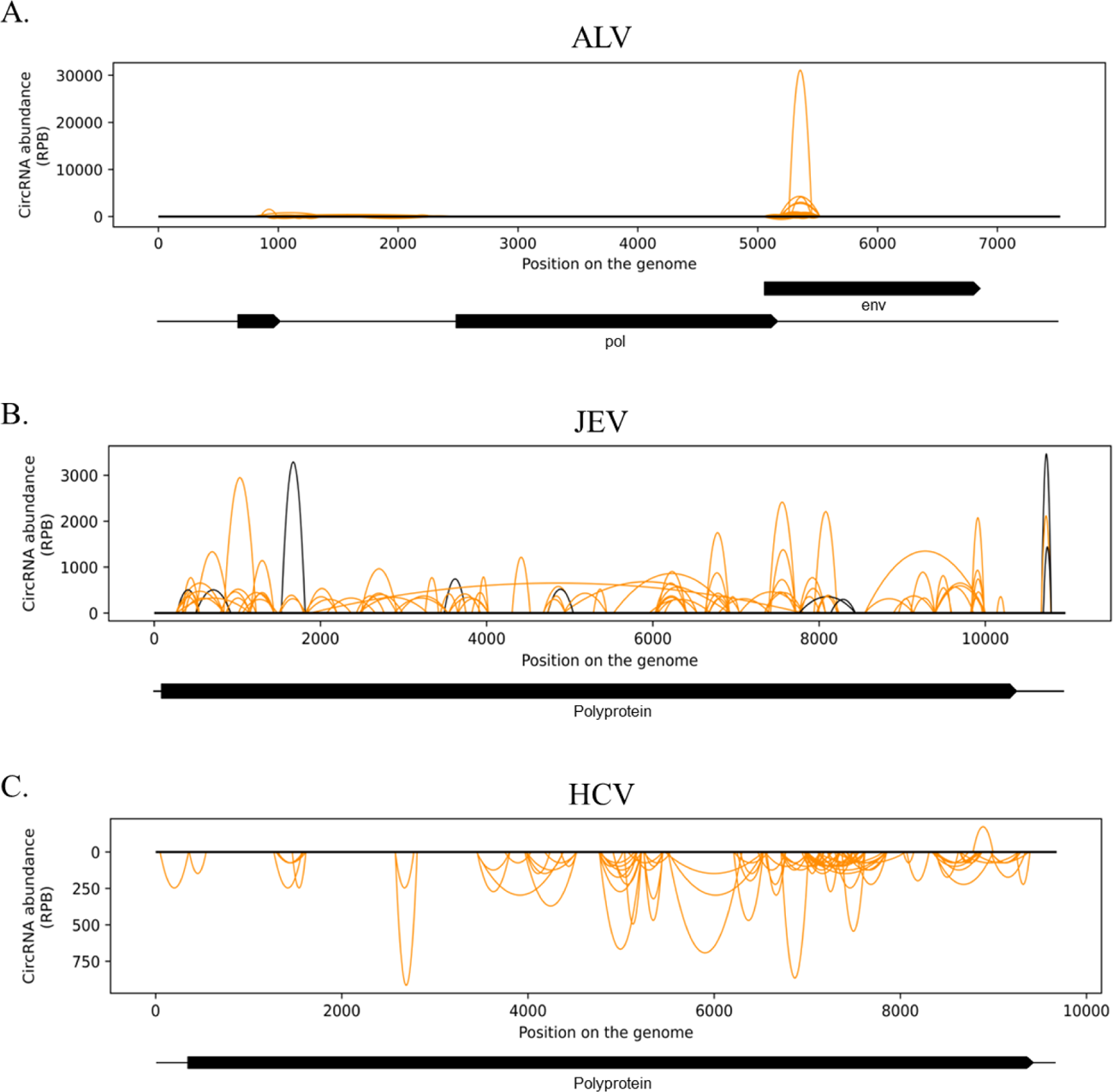
CircRNA identification in multiple viral infections. (**A**) Viral circRNAs identification from an ALV infection. The dataset was obtained after the infection of Chicken Embryonic Fibroblasts (CEF) in the study of Yang *et al*, 2022 (34). (**B**) Viral circRNAs identification from a JEV infection. Mouse brains were injected with the virus and harvested after 5 days of infection in the study of Li *et al*, 2020 (35). (**C**) Viral circRNAs identification from a HCV infection. Huh7 cells were infected with the JFH-1 strain of the virus in the context of the study of Cao *et al*, 2024 (11). The Y axis indicate the abundance of unique circRNAs in reads mapping the backsplice junctions per billion of reads mapping on the viral genome.

